# Auditory Working Memory Mediates the Relationship between Musical Sophistication and Speech-in-noise Perception

**DOI:** 10.64898/2026.05.13.724783

**Authors:** Hasan Colak, Ester Benzaquén, Xiaoxuan Guo, Meher Lad, William Sedley, Timothy D. Griffiths

## Abstract

Understanding speech in noisy environments (SPIN) is an important everyday ability, and engaging in musical activities has been proposed as a factor that may support this ability. However, the cognitive mechanisms underlying a potential musical advantage in SPIN perception remain unclear. Here we investigated whether musical sophistication is associated with better SPIN perception in a large population-based sample, and whether this relationship is mediated by auditory working memory (AWM), verbal working memory (VWM), or non-verbal intelligence. We recruited 203 participants and measured SPIN perception at both word and sentence levels. Musical sophistication was assessed using the Goldsmiths Musical Sophistication Index (Gold-MSI). AWM was measured using delayed matching of tone frequency or the modulation rate of amplitude modulated white noise, VWM was based on backward digit span task, and non-verbal intelligence used matrix reasoning. Mediation analyses revealed that AWM fully mediated the relationship between musical sophistication and SPIN perception, whereas VWM showed no mediation effect. Non-verbal intelligence showed a partial mediating effect. Additional control analyses using structural equation modelling revealed that the indirect effect through AWM remained significant after accounting for age, hearing thresholds, and non-verbal intelligence. Together, these findings suggest that individuals with greater musical sophistication demonstrate better daily life listening abilities, and that superior auditory working memory may be the key cognitive mechanism underlying this advantage.

## Introduction

One of the common challenges people face in everyday life is understanding speech in noisy environments, which requires a finely tuned interaction between peripheral hearing and auditory-cognitive mechanisms. About 42% of individuals report having difficulty perceiving speech-in-noise (SPIN) despite having normal hearing thresholds (Smith et al., 2019). SPIN difficulties can be a major health burden, as they may lead to social withdrawal and loneliness, and can be linked—directly or indirectly—to cognitive decline, ultimately affecting quality of life (Merten et al., 2022; Moore et al., 2014; Stam et al., 2016; Stevenson et al., 2022). Given these broad consequences, it is practically important to understand why some individuals cope well with speech in the presence of competing talkers while others struggle. Untangling the factors behind these differences, and understanding how they affect SPIN and to what extent, would be the first step towards developing strategies to manage SPIN difficulties.

One group that seems to have an advantage in understanding speech in noisy environments is people who engage in music, such as those who play an instrument or actively participate in musical activities like singing. There is growing evidence suggesting that musicians, or individuals who have undergone musical training for some period of time, outperform non-musicians in SPIN perception (Başkent & Gaudrain, 2016; Elisabeth Maillard et al., 2023; Morse-Fortier et al., 2017; Parbery-Clark et al., 2009; Parbery-Clark et al., 2011; Slater et al., 2015). (Boebinger et al., 2015; Madsen et al., 2019; Madsen, Whiteford, & Oxenham, 2017; Ruggles, Freyman, & Oxenham, 2014). One possible reason for these mixed findings is that different SPIN paradigms place different perceptual and cognitive demands on the listener. In particular, listening conditions in which speech is masked by competing speech may increase informational masking and place greater demands on higher-level processes such as selective attention and auditory stream segregation, which may be more sensitive to musical experience. It is also possible that musical training affects speech-on-speech processing in ways that do not always translate into better behavioural SPIN performance. For example, musicians may show earlier disambiguation of target speech from background, possibly due to training-related advantages in processes relevant to stream segregation (Kaplan et al., 2021). A meta-analysis by Hennessy, Mack and Habibi (2022) reported a consistent musician advantage in SPIN perception, irrespective of age, speech material, or IQ. Another line of evidence supporting this advantage comes from longitudinal studies showing that engaging in musical activities over time leads to improved SPIN perception (Dubinsky et al., 2019; E. Maillard et al., 2023; Slater et al., 2015). To explain the roots of this effect, the OPERA hypothesis (overlap, precision, emotion, repetition, and attention) proposes that shared auditory-cognitive mechanisms involved in processing music and speech are strengthened through musical training, depending on the degree of engagement (e.g., emotional reward, repetition, and attentional focus) (Patel, 2011, 2014). Specifically, speech and music perception are thought to rely on overlapping neural pathways and structures, from subcortical to cortical levels, where acoustic input is progressively processed and gain its meaning. Music processing also places particularly high demands on precision, as it requires fine-grained analysis of pitch, timing, and periodicity, all of which are also relevant to speech processing. In addition, musical activities are often emotionally rewarding, highly repetitive, and attentionally engaging, all of which may help drive plastic changes in the brain. Repetition gives listeners more chances to practise, attention helps them focus on the relevant sounds, and emotional reward may help keep them engaged and motivated to learn. The extent to which these shared mechanisms contribute to both speech and music processing is still being investigated, though several possible links have been proposed.

From first principles, the ability to group and maintain auditory elements over time is essential for forming auditory streams and achieving sound segregation. This ability enables both SPIN understanding and music processing by allowing us to sustain attention and retain relevant information in working memory. A large body of research on the role of working memory in SPIN perception suggests that, beyond peripheral hearing, the ability to hold relevant information in mind supports SPIN performance, such that the better one’s working memory, the better the understanding of SPIN (Benzaquén et al., 2025; Kim et al., 2020; Kraus, Strait, & Parbery-Clark, 2012; Lad et al., 2020; Rönnberg et al., 2013; Schulze & Koelsch, 2012; Yeend, Beach, & Sharma, 2019), but see Füllgrabe and Rosen (2016). Similarly, engaging in musical activities has been associated with enhanced working and short-term memory, particularly in the tonal domain (Grassi et al., 2025; Schulze & Koelsch, 2012; Talamini et al., 2017). These findings raise the question of whether the observed musical advantage in SPIN perception is driven by superior working memory. Zhang et al. (2021) reported a partial mediating role of verbal working memory, assessed with the digit span task, in the relationship between musical training and SPIN performance. However, this relationship may be more complex than it appears. Other higher level cognitive factors, such as non-verbal intelligence, may covary with working memory and could also account for this advantage. This interpretation resonates with previous work reporting higher non-verbal intelligence scores in musicians compared to non-musicians (Bidelman & Yoo, 2020; Schellenberg, 2011; Schellenberg & Lima, 2024; Yoo & Bidelman, 2019). One possible explanation for this difference is socioeconomic status, which has been linked to both participation in musical activities and higher intelligence (Albert, 2006; Bates, Lewis, & Weiss, 2013; Holster, 2023; von Stumm & Plomin, 2015). Boebinger et al. (2015) found no difference in SPIN perception between musicians and non-musicians when the groups were matched for non-verbal intelligence and working memory, and reported that SPIN performance was predicted by non-verbal intelligence rather than musical experience. Given this complex interplay of cognitive factors, further work is needed to better elucidate the origins of the musician advantage in real-world listening situations.

The majority of previous studies have focused primarily on professional musicians and therefore relied on relatively small sample sizes. Here, we collected data from a large population-based sample of individuals with varying levels of musical engagement including subjects with no musical training with varying degrees of engagement in music including singing and listening. We aimed to better understand whether greater musical sophistication is associated with better SPIN perception, and if so, whether this relationship is mediated by auditory or verbal working memory or non-verbal intelligence. We hypothesized that individuals with higher musical sophistication would show better SPIN performance, and that this advantage would be explained by superior working memory and possibly by higher non-verbal intelligence.

## Methods

### Participants

The present sample was drawn from the AudCog study, which focuses on higher-level auditory processing in cognitively healthy individuals and volunteers with cognitive disorders (Benzaquén et al., 2025). Recruitment was carried out through a range of channels, including local volunteer databases, institute participant databases, and public advertising. We analysed data from 203 participants (135 Female) aged between 18 and 81 years, all of whom had no diagnosis of cognitive or neurological disorder. All participants were native English speakers. Each participant underwent pure tone audiometry across the 0.25–8 kHz range using a calibrated clinical audiometer (AD629 Interacoustics), following the Modified Hughson–Westlake procedure, and thresholds were obtained in dB HL. Pure tone averages were calculated across frequencies from 0.25 to 8 kHz for both ears (PTAall), given the wide age range of the sample. This approach was used to account for higher frequencies, which are more sensitive to age-related hearing loss. Audiograms for all participants are shown in Figure 1. All testing took place in a soundproof and electrically shielded room, with participants seated comfortably. Custom computerized auditory tests were administered using Sennheiser HD 201 circumaural headphones, and stimuli were presented via Psychtoolbox in MATLAB 2017a or in Javascript. Practice trials were provided for all computerized tasks to familiarize participants with the procedures and stimuli. Additional details on the tasks can be found in Benzaquén et al. (2025). The study was approved by the Newcastle University Ethics Committee (Reference number: 46225/2023), and written informed consent was obtained from all participants.

**Figure 1.**
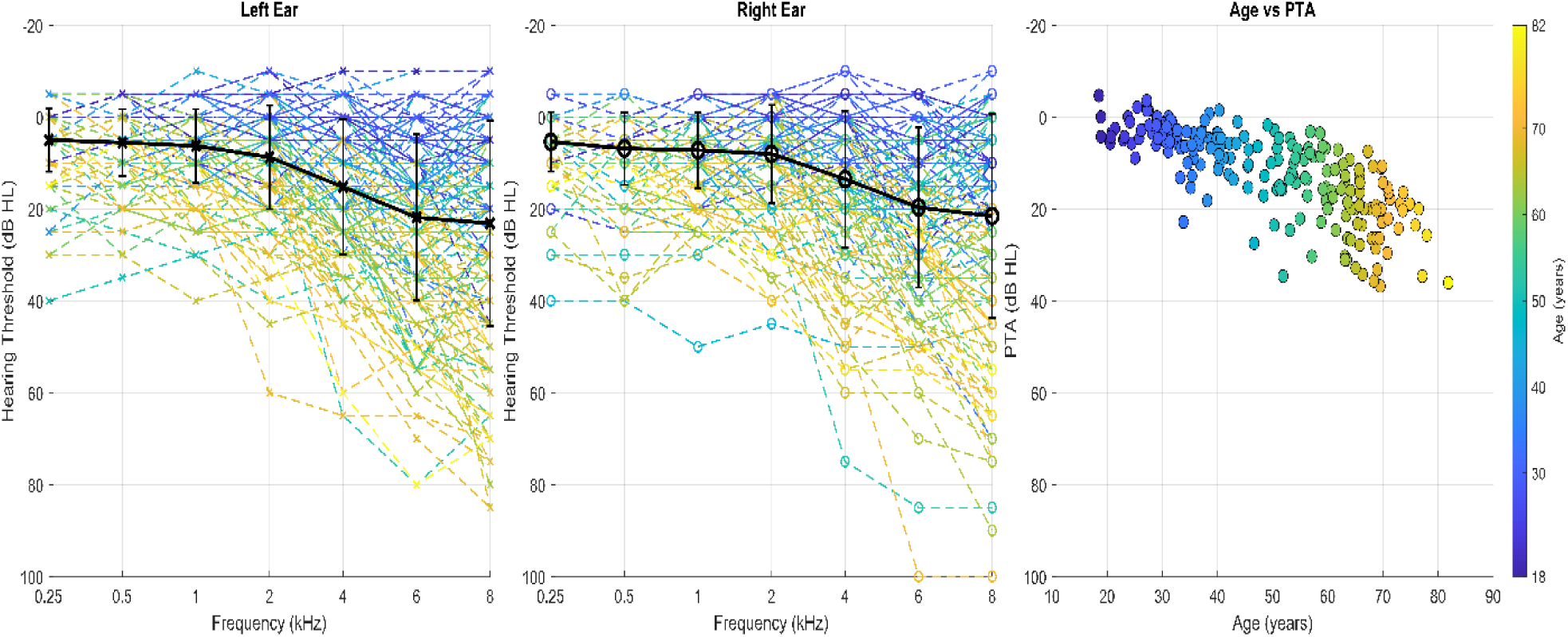
The audiometric thresholds of the participants. The left and middle panels show audiograms for the left and right ears, respectively. Thin dashed coloured lines represent individual participants, and thick black lines with error bars represent group means ± standard deviations across frequencies from 0.25 to 8 kHz. The right panel shows the relationship between age and pure-tone average across the frequencies (PTAall), with each point representing one participant and colour indicating age.

### Goldsmiths Musical Sophistication Index (Gold-MSI)

The Gold-MSI (Müllensiefen et al. (2014) is a self-report questionnaire that assesses musical skills, music-related behaviours, and engagement across several domains in the general population. It consists of 38 items distributed across five subscales: active engagement, perceptual abilities, musical training, singing ability, and emotional response to music. Each item is rated on a 1–7 Likert scale. The Gold-MSI allows musicality to be treated as a continuous variable, enabling the study of different levels of musical engagement rather than relying on a binary categorization (musician vs. non-musician).

### Working Memory

Working memory abilities were evaluated across two domains, auditory and verbal. The auditory working memory (AWM) task involved frequency and amplitude modulation (AM) precision, in which participants were required to remember a target sound across a silent delay and compare it with the different sounds they generated during the matching phase before selecting the best match (Lad et al., 2022; Lad et al., 2020). The target sound was either a 1-second pure tone or a 1-second sinusoidally amplitude-modulated white noise with 100% modulation depth, followed by a 2s silent delay after the target sound ended. For the matching of tone frequency, low-frequency stimuli ranging from 440 to 880 Hz were used to minimize potential confounds related to high-frequency hearing loss. For the matching of AM rate, the modulation rate varied from 5 to 20 Hz. The starting phase of the AM modulator was held constant across trials. Trial type was predetermined, such that participants matched either a pure tone or an AM noise depending on the target presented on that trial. After the delay period, participants were presented with a horizontal scale on which they could move a marker and click with the mouse to play their selected sound. The adjustment was continuous rather than step-based, and the position on the scale was mapped onto the relevant frequency or AM-rate range. They could listen and adjust without a time limit before finalizing their match to the target sound. Each participant completed a total of 32 trials, with 16 trials in each of the frequency and AM domains. The main outcome measure was precision, calculated as the inverse of the standard deviation of the matching errors for each domain. Matching error was calculated on a log scale as the difference between the target value and the participant’s selected value, relative to the target. All stimuli were presented at 70 dB SPL.

Verbal working memory (VWM) was assessed using the Digit Span Backward subtest from the Wechsler Memory Scale, Third Edition (WMS-III). In this task, digit sequences were presented orally by the experimenter, and participants were required to recall each sequence in reverse order and responded verbally. The test consisted of 7 items, each containing 2 trials, resulting in a maximum possible score of 14. Sequence length began at 2 digits in the first item and increased progressively to 8 digits in the seventh item. Testing was discontinued if a participant scored 0 on both trials of any item. Performance was scored as the total number of correctly recalled sequences.

### Non-verbal Intelligence

We used a matrix reasoning test (MTX) to assess the non-verbal domain of general intelligence. The matrices were taken from the Matrix Reasoning Item Bank (MaRs-IB; Chierchia et al. (2019)). The test included 30 trials in which participants were required to solve visual patterns by identifying the shape that correctly completed each matrix. The first four trials served as practice, and performance was scored as the total number of correctly solved matrices out of the remaining 26 trials. To prevent participants from spending excessive time on any single matrix, each trial was limited to 30 seconds, with a countdown timer displayed during the final 5 seconds.

### Speech-in-noise (SPIN) Perception

SPIN perception was tested at both the single-word and sentence levels. Single-word SPIN performance was measured using the British version of the Iowa Consonant Perception Test (Guo et al., 2024): a single-word closed-set speech-in-noise test with well-balanced phonetic features. The test included recordings from one female and one male speaker, both using a British Southern Standard accent. Trials from the two speakers were presented in a randomised order across the task. Participants completed 120 trials in which four phonetically balanced alternative words were displayed on the screen, and they were asked to select the target word. The target items were monosyllabic words with a consonant–vowel–consonant (CVC) structure (e.g., “ball–fall–shawl–wall”). Stimuli were presented at a fixed signal-to-noise ratio (SNR) of –2 dB, with the target words presented at 70 dB SPL against an 8-talker babble background, consistent with the validated protocol and intended to better reflect everyday listening environments with multiple competing talkers, such as cafés, classrooms, or other busy social settings. Performance was quantified as the percentage of correctly identified target words out of 120 trials.

The sentence-in-noise (SiN) test was implemented in JavaScript using the English version of the Oldenburg Sentence Test (Holmes & Griffiths, 2019). The target sentences followed a fixed five-word structure: <Name> <Verb> <Number> <Adjective> <Noun>. Each sentence was presented against a 16-talker babble background, as in the original study (Holmes & Griffiths, 2019). Participants were shown a 5 × 10 response matrix on the screen, which contained 10 word options for each word category, and were asked to select the words they heard. The test followed an adaptive 1-up 1-down procedure, beginning at a signal-to-noise ratio (SNR) of 10 dB, with all stimuli presented at 70 dB SPL. The target speech level remained constant, while the noise level was adjusted based on participant performance. Step sizes decreased from 3 to 2 dB after three reversals, and further to 1 dB after three additional reversals. For the final six reversals, the step size was reduced to 0.5 dB. Each run terminated after 12 reversals, and the median SNR of the last six reversals was taken as the threshold score for each participant. Lower SNR thresholds indicated better performance.

## Statistical Analysis

Statistical analyses were conducted in MATLAB R2024a and in RStudio. Descriptive statistics for all variables are reported in Table 1. Prior to analysis, all continuous measures were z-scored to place them on a common scale, as the original variables were measured in different units. Overall SPIN performance was calculated by summing the z-scored word-in-noise (WiN) and sentence-in-noise (SiN) measures, after reversing the direction of the SiN scores so that higher values consistently reflected better performance. Similarly, overall AWM performance was calculated by summing the z-scored frequency and AM AWM measures. Spearman correlation coefficients were computed to examine the bivariate associations among total Gold-MSI score, AWM, VWM, MTX, SPIN performance, PTA, and age. Effect sizes were interpreted based on Spearman’s rho (ρ), where 0.10 indicated a small effect, 0.30 a medium effect, and 0.50 a large effect. The p-values were adjusted for multiple comparisons using the Benjamini–Hochberg false discovery rate procedure. Multicollinearity was assessed by calculating variance inflation factor (VIF) values for all variables. All VIF values were below 2, indicating that collinearity was not a concern.

**Table 1.** Descriptive statistics of the participants. Means, standard deviations, minimum values, and maximum values are reported for age, pure-tone average across 0.25–8 kHz for both ears (PTAall), musical sophistication as measured by the Goldsmiths Musical Sophistication Index (Gold-MSI), auditory working memory (AWM; frequency and amplitude modulation subtests), verbal working memory (VWM), non-verbal intelligence (MTX), and speech-in-noise performance on the word-in-noise (WiN) and sentence-in-noise (SiN) tasks. P.E. = precision estimate; SNR = signal-to-noise ratio; HL = hearing level.

|  | Mean | Standard Deviation | Min | Max |
| --- | --- | --- | --- | --- |
| Age (years) | 49.15 | 16.13 | 18.49 | 81.85 |
| PTAall (dB HL) | 12.01 | 9.35 | -4.64 | 36.78 |
| Gold-MSI (total score) | 145.03 | 33.63 | 51 | 230 |
| AWM Frequency (P.E) | 2.11 | 1.53 | 0.28 | 6.48 |
| AWM Amplitude Modulation (P.E) | 0.21 | 0.08 | 0.05 | 0.55 |
| VWM (total correct) | 7.31 | 2.14 | 4 | 13 |
| MTX (total correct) | 14.29 | 4.32 | 3 | 25 |
| WiN (%) | 66.67 | 8.89 | 44.16 | 87.5 |
| SiN (dB SNR) | -2.23 | 1.68 | -6 | 2.75 |

To better understand the mechanistic relationship between musicality and SPIN perception, we constructed several mediation models. In the mediation analyses, path coefficients were estimated using the ‘fitlm’ function, and indirect effects were tested using nonparametric bootstrapping implemented with the ‘bootstrp’ function. In the first model, AWM was tested as a mediator of the association between musicality and SPIN performance. In the second model, VWM was examined as an alternative mediator to determine whether verbal working memory explains additional variance in SPIN outcomes. In the third model, MTX was tested as another potential mediator. For each model, path coefficients (a, b, c, c′) were estimated using linear regression. Indirect effects were calculated as the product of the a and b paths. Significance was assessed using nonparametric bootstrapping with 5,000 resamples. The bootstrap distribution was used to compute bias-corrected 95% confidence intervals and empirical two-tailed p-values.

Finally, we conducted two structural equation modelling (SEM) analyses in RStudio using the lavaan package as control analyses to test whether the indirect effect of musicality through AWM remained significant after accounting for individual differences in non-verbal intelligence (MTX), hearing thresholds (PTAall), and age. Models were estimated using maximum likelihood with correction for non-normality based on the Satorra–Bentler scaled test statistic. Model fit was assessed using several goodness-of-fit indices: the comparative fit index (CFI), Tucker–Lewis index (TLI), root-mean-square error of approximation (RMSEA). Following common guidelines, CFI and TLI values around 0.90 or above were taken to indicate acceptable fit, values around 0.95 or above good fit, and RMSEA values around 0.08 or below acceptable fit, values around 0.06 or below good fit. Only robust versions of these indices are reported. Bootstrapped 95% confidence intervals for model fit measures were generated using 1000 repetitions with the Bollen–Stine method. The structural and latent components of the models were specified in line with theoretical expectations.In the first model, SPIN and AWM were specified as latent variables, where SPIN was indicated by WiN and SiN performance and AWM by the frequency and AM auditory working memory measures. In the second model, we specified a broader working memory latent factor, indicated by the frequency and AM auditory working memory measures together with verbal working memory performance, to test whether the mediation effect remained when verbal working memory was incorporated into the latent construct. In both models, the indirect effect of musicality on SPIN was tested while including MTX, PTAall, and age as potential confounds.

## Results

### Participant Characteristics

The mean age of the participants was 49.15 years (SD = 16.13). The mean pure tone average (0.25–8 kHz) was 12.01 dB HL (SD = 9.35; see Figure 1). Additional descriptive statistics for all measures are presented in Table 1.

### Relationship Between Musicality and SPIN Performance

The bivariate analysis showed that musicality was significantly associated with SPIN performance, with a small effect size, even after correction for multiple comparisons (ρ= 0.26, corrected p < 0.001; see Figure 2). When examined separately, musicality was significantly correlated with both word-level and sentence-level SPIN performance (WiN: ρ = 0.25, corrected p < 0.001; SiN: ρ = - 0.22, corrected p = 0.002).

**Figure 2.**
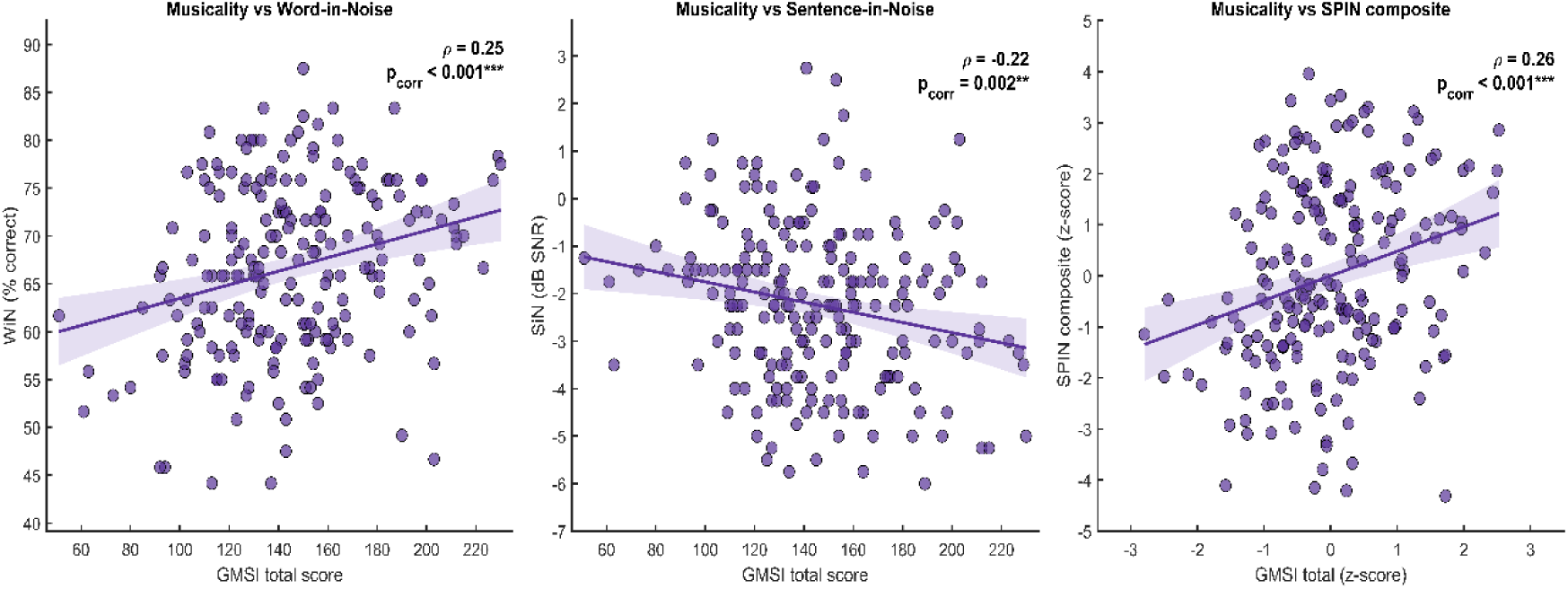
The association between musicality and SPIN perception. Scatterplots showing the associations between musicality and speech-in-noise performance. The left panel shows the relationship between total Goldsmiths Musical Sophistication Index (Gold-MSI) score and word-in-noise (WiN) performance, expressed as percentage correct. The middle panel shows the relationship between total Gold-MSI score and sentence-in-noise (SiN) performance, expressed in dB SNR, where lower values indicate better performance. The right panel shows the relationship between z-scored Gold-MSI total score and the composite speech-in-noise (SPIN) score, calculated by z-scoring the WiN and SiN measures and summing them after reversing the direction of the SiN scores so that higher values reflected better performance. Each point represents one participant. Solid lines represent linear fits, and shaded areas indicate 95% confidence intervals. Spearman’s rho and corrected p values are shown in the upper right corner of each panel.

### Cognitive Correlates of SPIN performance and Musicality

We further examined whether the cognitive variables were associated with SPIN performance and musicality. SPIN performance showed significant positive associations with AWM (ρ = 0.48, corrected p < 0.001), VWM (ρ = 0.22, corrected p = 0.001), and MTX (ρ = 0.42, corrected p < 0.001). Musicality was also significantly correlated with both AWM (ρ = 0.44, corrected p < 0.001) and MTX (ρ = 0.23, corrected p = 0.001). In contrast, musicality was not associated with VWM (ρ = 0.005, corrected p = 0.938). The correlation results are presented in Table 2.

**Table 2.** Spearman correlations between SPIN performance, musicality, cognitive measures, PTAall, and age.

|  | SPIN | WiN | SiN | Musicality | PTAall | Age |
| --- | --- | --- | --- | --- | --- | --- |
|  | (WiN+SiN) |  |  | (Gold-MSI) |  |  |
| AWM | $\rho = 0.48$ | $\rho = 0.48$ | $\rho = -0.38$ | $\rho = 0.44$ | $\rho = -0.37$ | $\rho = -0.34$ |
| | $p < 0.001$ | $p < 0.001$ | $p < 0.001$ | $p < 0.001$ | $p < 0.001$ | $p < 0.001$ |
| VWM | $\rho = 0.22$ | $\rho = 0.20$ | $\rho = -0.18$ | $\rho = 0.005$ | $\rho = -0.06$ | $\rho = 0.01$ |
| | $p = 0.001$ | $p = 0.004$ | $p = 0.008$ | $p = 0.938$ | $p = 0.402$ | $p = 0.845$ |
| MTX | $\rho = 0.42$ | $\rho = 0.44$ | $\rho = -0.33$ | $\rho = 0.23$ | $\rho = -0.37$ | $\rho = -0.36$ |
| | $p < 0.001$ | $p < 0.001$ | $p < 0.001$ | $p = 0.001$ | $p < 0.001$ | $p < 0.001$ |

Spearman correlation coefficients ( $\rho$ ) between auditory working memory (AWM), verbal working memory (VWM), non-verbal intelligence (MTX), speech-in-noise (SPIN) performance, word-in-noise (WiN), sentence-in-noise (SiN), musicality (Gold-MSI), pure tone average (PTAall), and age. SPIN performance reflects a composite score derived from WiN and SiN measures. All p-values were adjusted for multiple comparisons using the Benjamini–Hochberg false discovery rate procedure.

### Mediation Models

Given the significant correlation between musicality and SPIN performance, we conducted mediation analyses to clarify the mechanisms underlying this relationship and to assess the potential roles of working memory and non-verbal intelligence. In the first model, AWM was tested as a mediator of the association between musicality and SPIN perception. Results indicated a strong and significant effect of musicality on AWM (path a = 0.80, p < 0.001), and AWM significantly predicted SPIN performance when controlling for musicality (path b = 0.45, p < 0.001). The direct effect of musicality on SPIN performance was not significant once AWM was included in the model (path c′: β = 0.11, p = 0.38), whereas the total effect was significant (path c: β = 0.48, p < 0.001). The indirect effect was statistically significant (0.37, p < 0.001). Together, these results indicate a full mediation, suggesting that AWM fully accounts for the relationship between musicality and SPIN performance (Figure 3).

**Figure 3.**
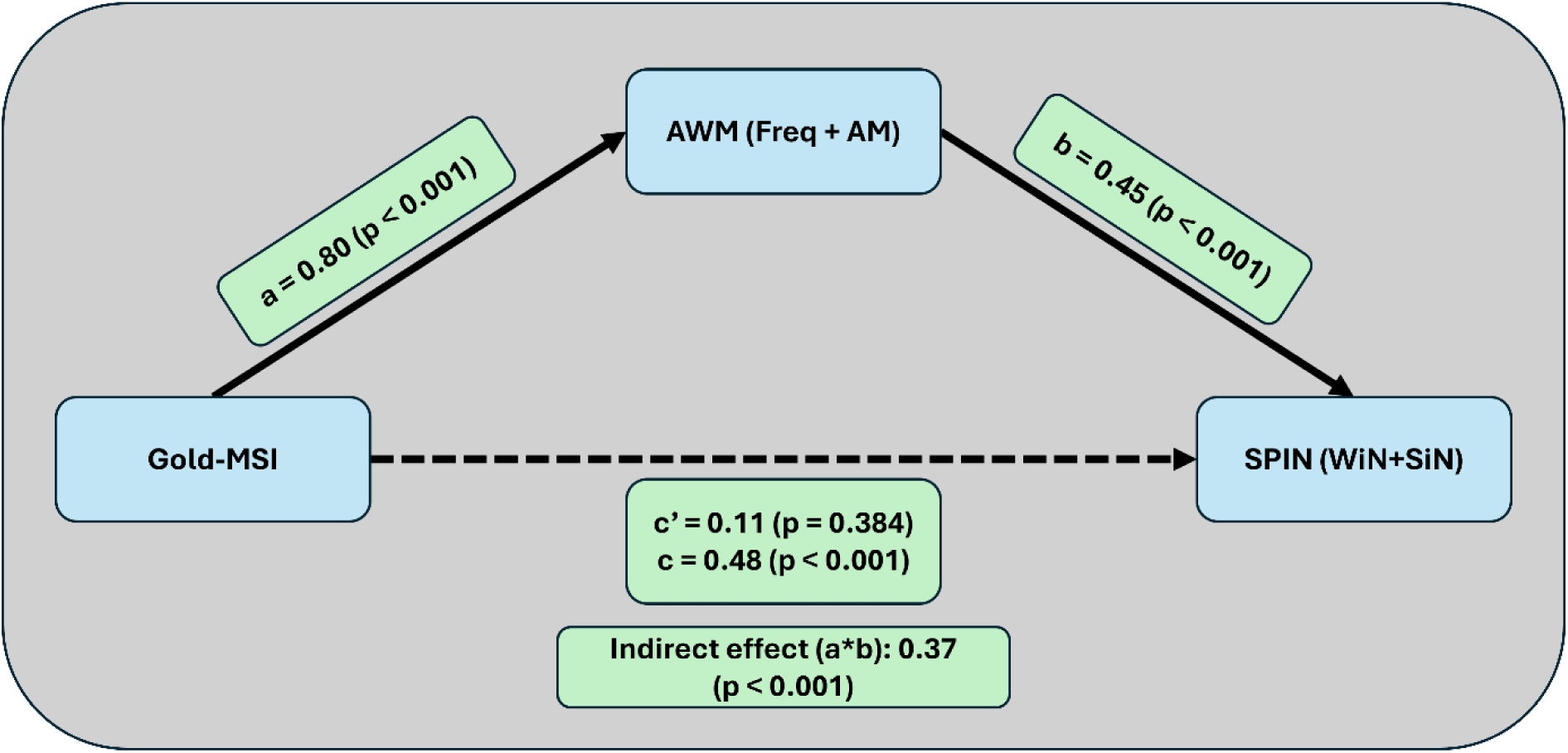
Mediation model showing the indirect effect of musicality on SPIN performance through AWM. Gold-MSI total score was used as the measure of musicality. AWM was derived by combining the frequency and amplitude modulation (AM) subtests, and SPIN was derived by combining word-in-noise (WiN) and sentence-in-noise (SiN) performance. Path a represents the relationship between musicality and AWM. Path b reflects the relationship between AWM and SPIN performance. Path a × b illustrates the indirect effect of musicality on SPIN performance through AWM. Path c′ indicates the direct relationship between musicality and SPIN performance after accounting for the mediating role of AWM, and path c represents the total effect of musicality on SPIN performance.

We next examined whether VWM played a similar mediating role to AWM in the relationship between musicality and SPIN performance. Musicality did not significantly predict VWM (path a = 0.02, p = 0.75), indicating no meaningful association between the two measures. Although VWM significantly predicted SPIN performance when controlling for musicality (path b = 0.42, p < 0.001), the direct effect of musicality on SPIN performance remained strong and significant (path c′: β = 0.47, p < 0.001). Consistent with this pattern, the indirect effect was very small (0.009) and non-significant (p = 0.74). These results indicate no evidence of mediation, suggesting that VWM does not account for the relationship between musicality and SPIN performance (Figure 4).

**Figure 4.**
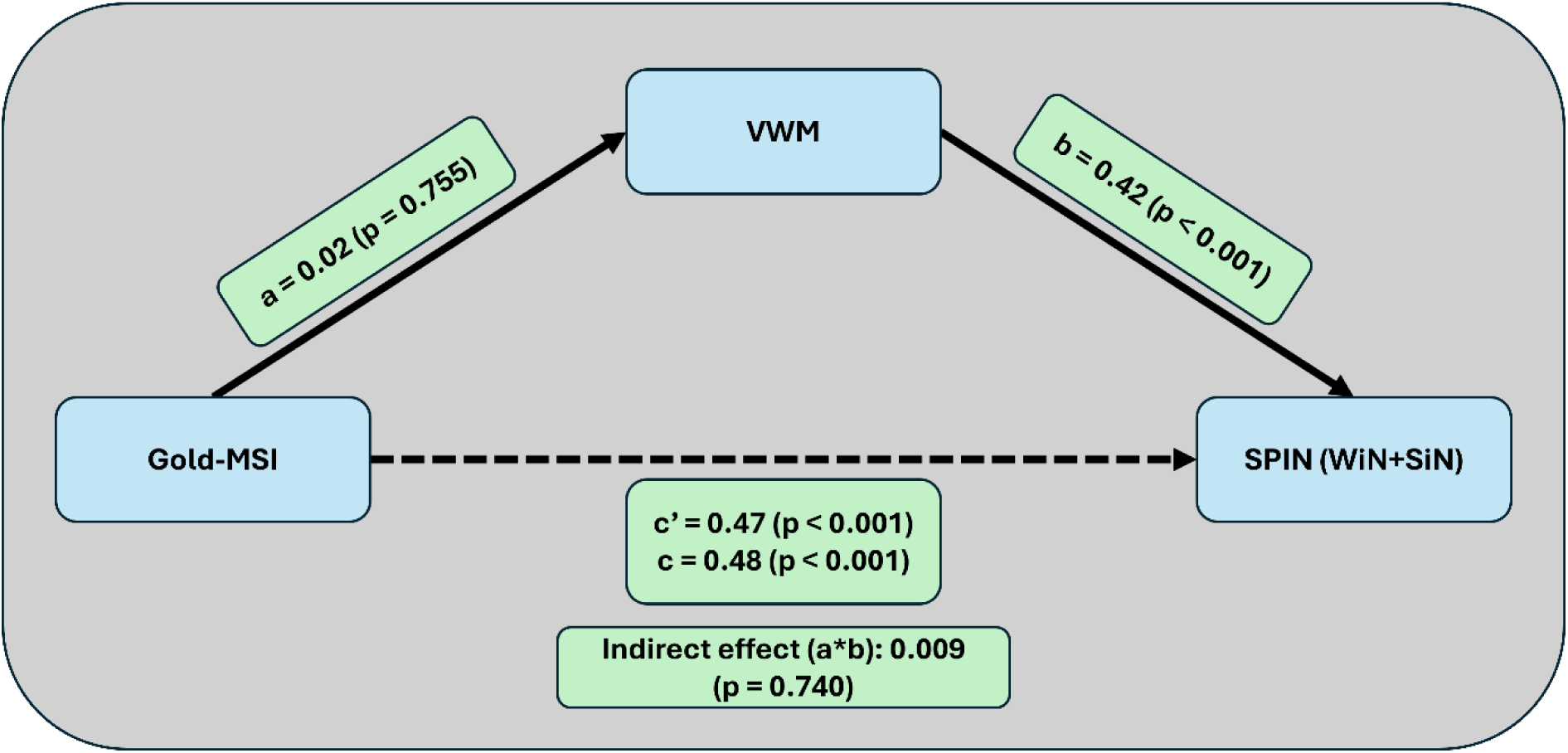
Mediation Model with VWM as Mediator. Gold-MSI total score was used as the measure of musicality, verbal working memory (VWM) was indexed using the total score on the backward digit span task, and SPIN was derived by combining word-in-noise (WiN) and sentence-in-noise (SiN) performance. Path a represents the relationship between musicality and VWM. Path b reflects the relationship between VWM and speech-in-noise (SPIN) performance. Path a × b illustrates the indirect effect of musicality on SPIN performance through VWM. Path c′ indicates the direct relationship between musicality and SPIN performance after accounting for the mediating role of VWM, and path c represents the total effect of musicality on SPIN performance

We also tested whether non-verbal intelligence mediated the association between musicality and SPIN performance. Musicality significantly predicted MTX (path a = 0.22, p = 0.001), and MTX showed a strong positive association with SPIN performance when controlling for musicality (path b = 0.71, p < 0.001). The direct effect of musicality on SPIN performance remained significant, though reduced in magnitude, when MTX was included in the model (path c′: β = 0.32, p = 0.007). The indirect effect was significant (0.16, p < 0.001). Taken together, these findings indicate that MTX partially mediates the relationship between musicality and SPIN performance (Figure 5).

**Figure 5.**
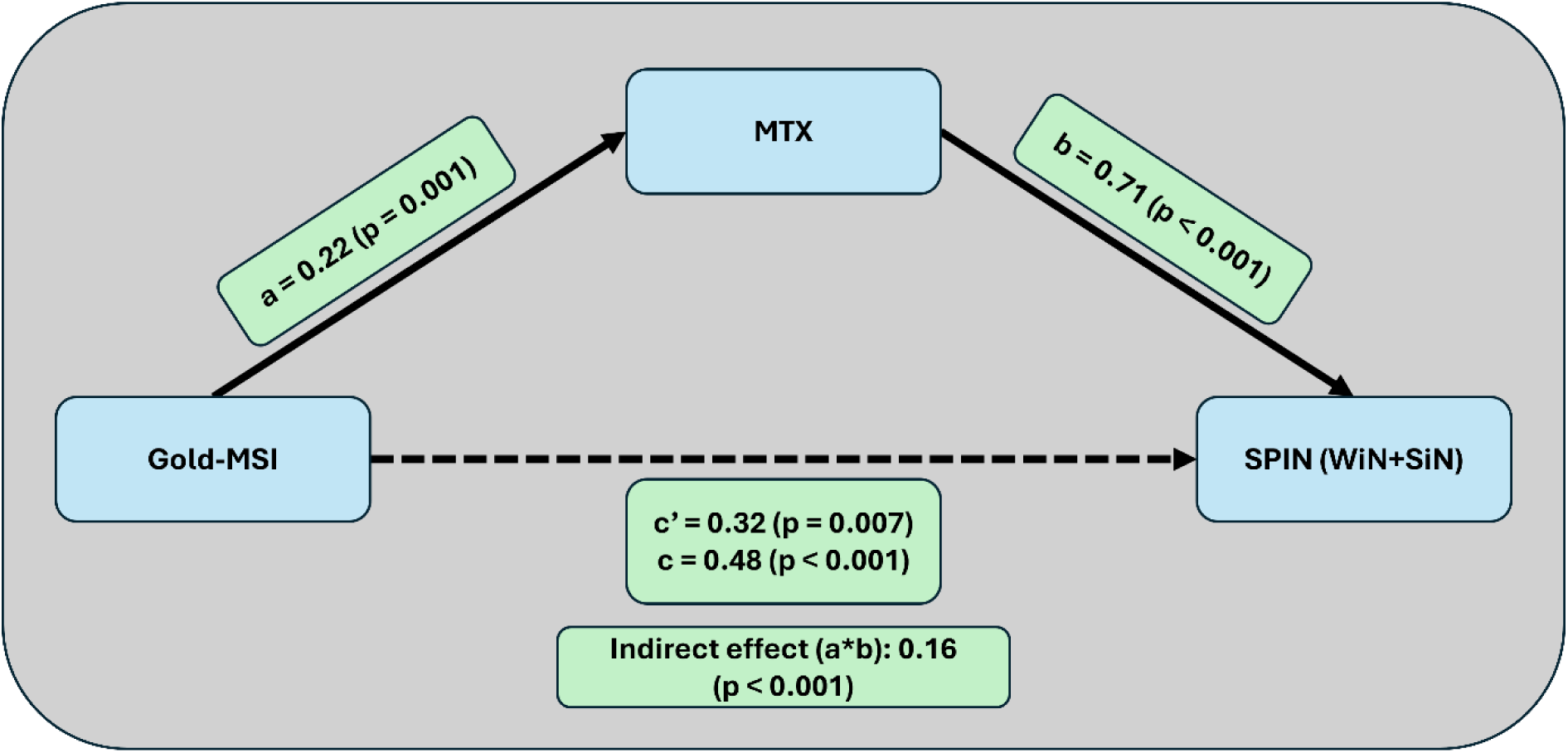
Mediation Model Testing the Role of Non-verbal Intelligence in the Relationship between Musicality and SPIN perception. Gold-MSI total score was used as the measure of musicality, MTX refers to performance on the matrix reasoning task, and SPIN was derived by combining word-in-noise (WiN) and sentence-in-noise (SiN) performance. Path a shows the association between musicality and MTX. Path b shows the relationship between MTX and SPIN perception. The path a × b term represents the indirect effect of musicality on SPIN performance transmitted through MTX. Path c′ reflects the direct effect of musicality on SPIN performance after accounting for MTX, while path c corresponds to the total effect of musicality on SPIN performance.

### Secondary Analyses

To test whether the main mediation finding remained after accounting for non-verbal intelligence, hearing thresholds (PTAall), and age, we conducted two SEM analyses as control analyses. In the first model, SPIN and AWM were specified as latent variables. Musicality significantly predicted latent AWM (β = 0.51, p < 0.001), and latent AWM significantly predicted latent SPIN (β = 0.28, p = 0.002). The overall effect of musicality on SPIN, calculated as the sum of the direct and indirect effects, was significant (β = 0.10, p = 0.016). However, the direct path from musicality to SPIN was not significant after accounting for AWM and the other confounds (β =-0.04, p = 0.579), whereas the indirect effect remained significant (β = 0.15, p = 0.014). Model fit indices for this SEM were CFI = 0.968, TLI = 0.931, and RMSEA = 0.093, indicating acceptable to good fit overall, although the RMSEA suggested only mediocre fit. This model is shown in Figure 6.

**Figure 6.**
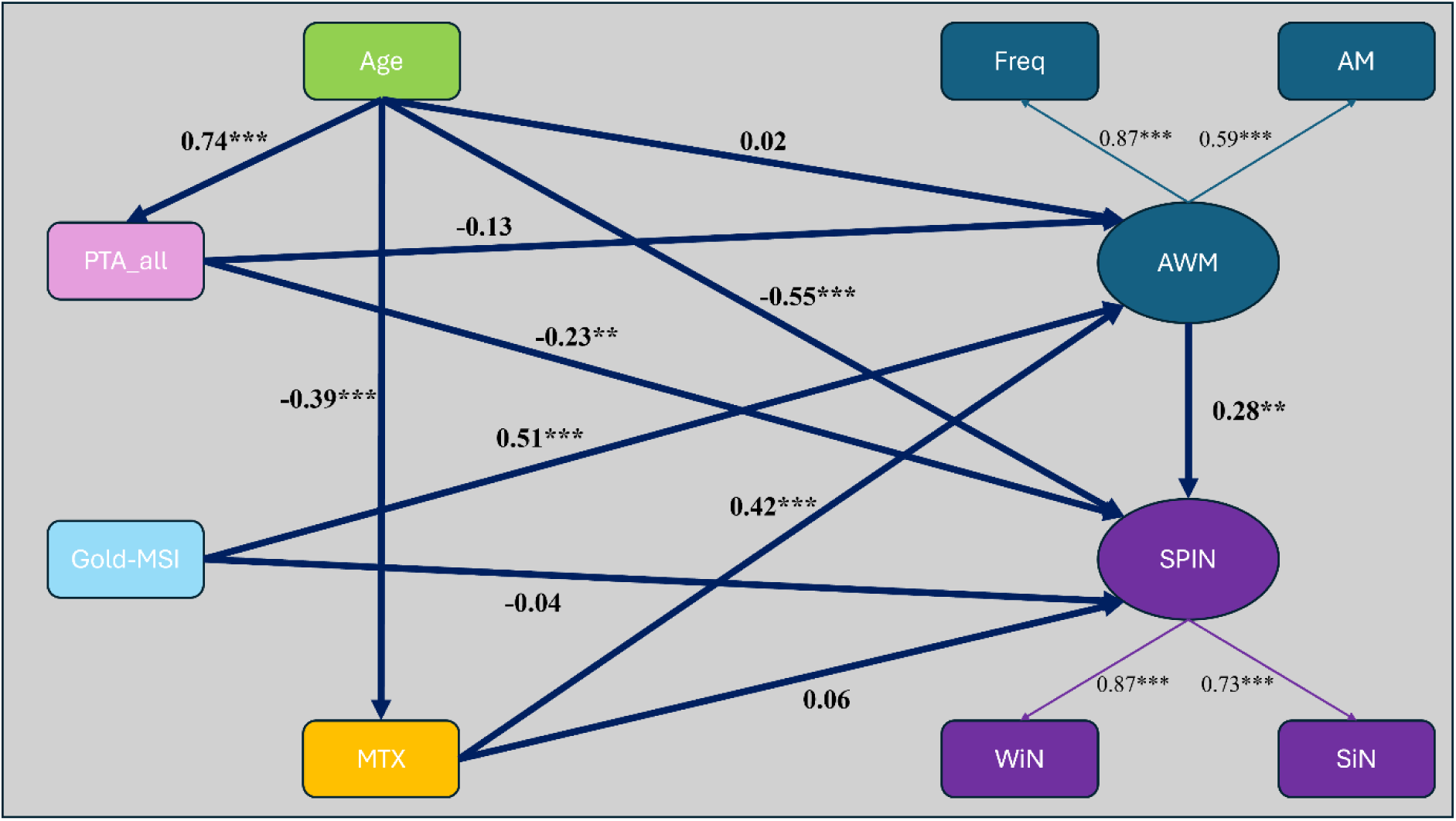
Structural equation model testing whether the indirect association between musicality and SPIN performance through AWM remained significant after accounting for age, hearing thresholds, and non-verbal intelligence. SPIN was modelled as a latent variable indicated by word-in-noise (WiN) and sentence-in-noise (SiN) performance, and AWM as a latent variable indicated by frequency (Freq) and amplitude modulation (AM) auditory working memory measures. Musicality was indexed using total Goldsmiths Musical Sophistication Index (Gold-MSI) score. Pure tone average calculated across 0.25–8 kHz for both ears (PTAall), and non-verbal intelligence was indexed using matrix reasoning (MTX). Values shown are standardised path coefficients. Asterisks indicate significance levels (p < 0.05*, p < 0.01**, p < 0.001***).

In the second SEM, we specified a broader latent working memory factor by combining the frequency and AM auditory working memory measures with verbal working memory (backward digit span) performance. This second model showed a similar pattern, in which musicality significantly predicted the working memory latent factor (β = 0.49, p < 0.001), which in turn significantly predicted latent SPIN (β = 0.32, p = 0.001). Again, total effect of musicality on SPIN was significant (β = 0.10, p = 0.016). The direct path from musicality to SPIN was not significant (β =-0.05, p = 0.471), whereas the indirect effect remained significant (β = 0.16, p = 0.008). Model fit indices for this second SEM were CFI = 0.946, TLI = 0.902, RMSEA = 0.100, indicating acceptable fit overall, although the RMSEA was relatively high. The corresponding SEM is shown in Figure 7.

**Figure 7.**
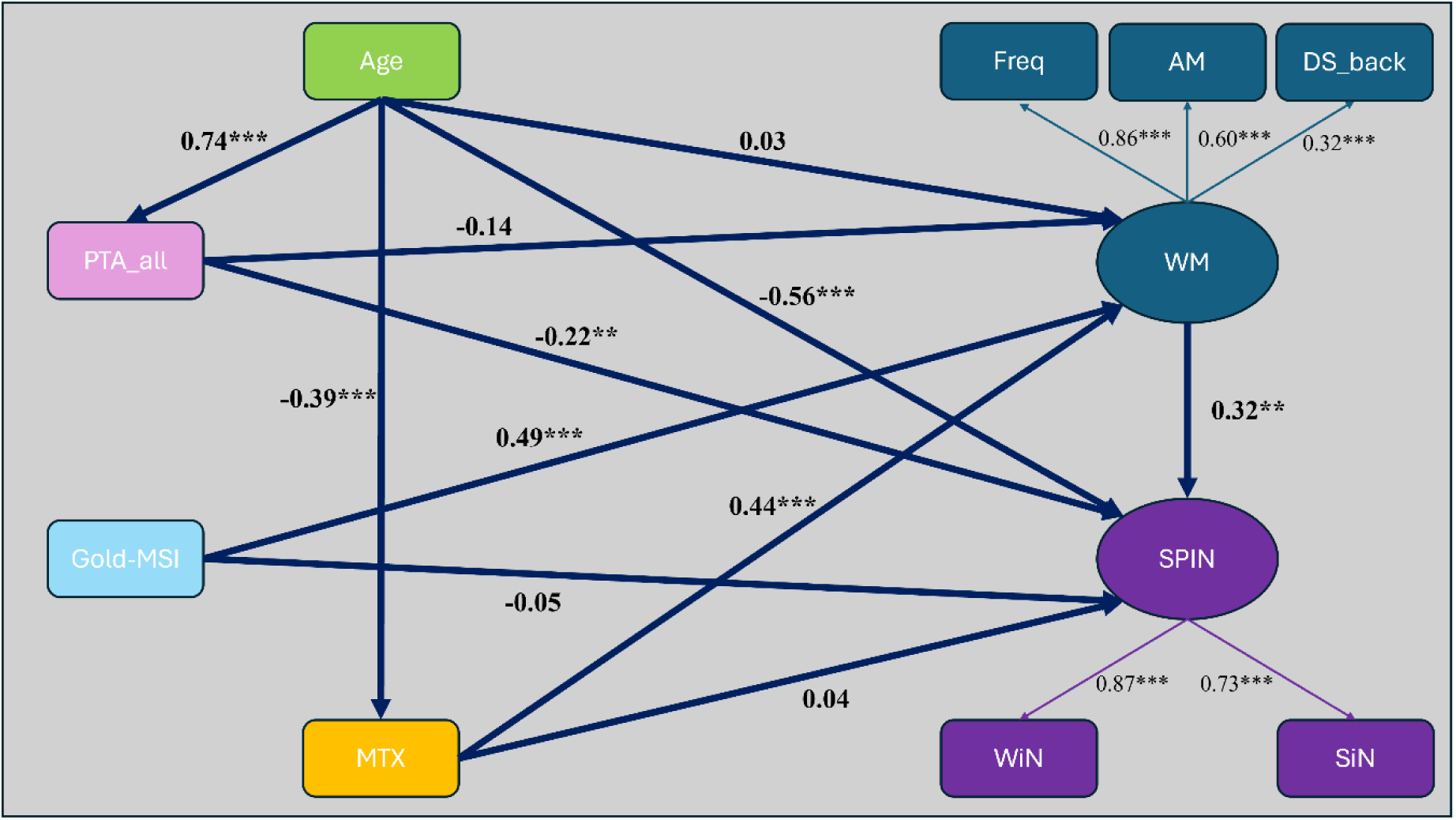
Structural equation model illustrating the indirect association between musicality and SPIN performance through a working memory (WM) latent factor, while accounting for age, hearing thresholds, and non-verbal intelligence. WM was modelled as a latent variable indicated by auditory working memory frequency (Freq), auditory working memory amplitude modulation (AM), and backward digit span performance (DS_back). SPIN was modelled as a latent variable indicated by word-in-noise (WiN) and sentence-in-noise (SiN) performance. Musicality was indexed using total Goldsmiths Musical Sophistication Index (Gold-MSI) score, hearing thresholds using pure tone average across 0.25–8 kHz for both ears (PTAall), and non-verbal intelligence using matrix reasoning (MTX). Values shown are standardised path coefficients. Asterisks indicate significance levels (p < 0.05*, p < 0.01**, p < 0.001***).

Together, these control analyses suggest that the association between musicality and SPIN is primarily indirect and remains evident after accounting for non-verbal intelligence, hearing thresholds, and age, including when verbal working memory is incorporated into the latent working memory construct.

## Discussion

Here we studied a large, population-based sample to address whether musical sophistication is associated with better SPIN perception and to identify the roles of working memory and non-verbal intelligence. Musicality was significantly associated with SPIN performance, such that higher musical sophistication was linked to better SPIN perception. This significant relationship was fully mediated by AWM, meaning that AWM ability drives this association. Interestingly, the results indicated that VWM did not show any mediation effect in the relationship between musicality and SPIN perception. Non-verbal intelligence also partially mediated this relationship, suggesting that it has a role in the association, albeit partial. Importantly, additional control analyses using SEM to account for potential confounds, including hearing thresholds, age, and non-verbal intelligence, showed that the indirect effect through AWM remained significant. A similar result was observed when verbal working memory was incorporated into a working memory latent factor. Together, these findings suggest that the association between musicality and SPIN is strongly linked to auditory working memory, while also indicating that this relationship is not fully explained by non-verbal intelligence, hearing, or age.

Although we did not specifically study trained musicians, our findings indicate that greater musical sophistication is associated with an advantage for everyday speech-in-noise abilities in the general population. Many previous studies on the musician advantage mostly rely on relatively small sample sizes and rarely account for intelligence, despite evidence that intelligence is associated with both musical training (Schellenberg, 2011) and SPIN perception (Dryden et al., 2017; Marsja et al., 2025; Moberly et al., 2018). Our mediation model with non-verbal intelligence as a potential mediator of the link between musicality and SPIN perception revealed a significant partial mediation effect.

### Non-verbal Auditory Working Memory fully mediates SPIN

An important aspect of the current work is that we examined whether the link between musicality and SPIN abilities is driven by superior working memory. Our findings indicate that AWM fully mediates the relationship between musicality and SPIN perception, and importantly, this effect remained significant in the additional SEM control analyses that accounted for hearing thresholds, age, and non-verbal intelligence. Previous work using the same AWM task and the Gold-MSI reported that musicality is not related to the perceptual components of the AWM task, such as frequency or AM discrimination (Lad et al., 2022). This supports the idea that AWM may be the cognitive factor linking musical sophistication to improved SPIN abilities in the general population. A recent study by Liu et al. (2025) showed that AWM, measured using tonal stimuli, mediates the relationship between musical training and auditory sound segregation. Our findings extend this effect to everyday listening and demonstrate that the same pattern applies to SPIN perception. Interestingly, VWM did not significantly mediate the relationship between musicality and SPIN perception. The larger musician advantage previously reported for working memory tasks that use tonal rather than verbal stimuli is consistent with our finding of a significant mediation effect for the AWM task (Schulze & Koelsch, 2012; Talamini et al., 2017; Yurgil et al., 2020). However, the lack of a mediation effect for VWM in our study contrasts with Zhang et al. (2021), who reported that VWM measured with digit span partially mediated the effect of musical training on SPIN performance. One possible explanation for this difference is that our sample consisted of more general population, whereas their study focused specifically on trained musicians. These differences in sample characteristics may underlie the seemingly contradictory findings.

One might still expect this superior working memory in the non-verbal auditory domain to extend to verbal domain as well, but this was not the case in our study. A key difference between the working memory tasks used here, in addition to the stimulus domain, is that the backward digit span task used to assess VWM requires participants to recall sequences of digits and mentally reverse their order. This introduces a manipulation requirement that engages central executive processes. This type of reordering and mental manipulation has been proposed to involve executive functioning (Hester, Kinsella, & Ong, 2004; Jahanshahi et al., 2008; Rosenthal et al., 2006). The AWM task we used places much lower demands on the central executive. It does not require manipulation or rehearsal of the stimuli and instead focuses on maintaining auditory information while selecting the correct match. Although some executive control is needed for the response, the absence of a manipulation component makes the AWM task less demanding than the backward digit span task. Although there is evidence that musical training can improve executive functions, particularly in younger populations such as children (Jamey et al., 2024; Román-Caballero et al., 2022), these additional central executive demands in the backward digit span may be less sensitive to plastic changes in adulthood, or may not transfer as readily through musical sophistication in a more general population. It is also possible that any effect of musical sophistication is less likely to translate as directly to the verbal domain as to the non-verbal auditory domain, particularly when the latter involves tones and AM sounds that are more closely related to the acoustic structure of music. This would be consistent with the fact that music relies heavily on pitch and rhythm, which map more directly onto these non-verbal auditory dimensions. As a result, any effects that do extend to the verbal domain may be smaller and therefore harder to detect in a population-based sample such as the one studied here. The absence of a mediation effect for VWM may therefore reflect both the different task demands and the more limited transfer of musical sophistication to verbal working memory at the population level.

### Non-verbal intelligence partially mediates SPIN

The reason why non-verbal intelligence supports SPIN perception is still unclear, but the two may overlap at certain cognitive levels. One possibility is that the brain processes involved in non-verbal intelligence tasks are similar to those required for auditory closure, where listeners must piece together incomplete auditory information. The significant association between musicality and non-verbal intelligence found here is also noteworthy. Although musicians are sometimes reported to have higher intelligence scores, this is less likely to be a direct result of musical training. Rather, factors such as socioeconomic background may contribute to both greater musical sophistication and higher intelligence. Taken together, our findings suggest that higher non-verbal intelligence explains part of the SPIN advantage associated with higher musical sophistication. This mediating role indicates that non-verbal intelligence should be considered a potential confound when interpreting musical advantages. It may also help explain why musicians did not differ from non-musicians in SPIN perception when they were matched on non-verbal intelligence (Boebinger et al., 2015). Future work on musical advantages in SPIN should therefore take non-verbal intelligence into account as an important covariate.

### Further considerations

There are several limitations of the current work that should be addressed in future studies. First, although the Gold-MSI correlates well with musical abilities such as melody and beat perception (Müllensiefen et al., 2014), it is a self-report questionnaire and may not provide direct evidence of the one’s actual musical ability. Future studies should therefore incorporate behavioural measures that assess different aspects of musical perception (e.g., melody, rhythm, timbre). Second, although previous work has shown that the AWM task used here is not related to perceptual discrimination of tone frequency or AM sounds (Lad et al., 2022), we were unable to verify this directly in the current dataset. Future work would benefit from more clearly separating the memory component of the task from the perceptual properties of the stimuli to ensure that the AWM measure is not influenced by basic auditory discrimination abilities.

## Conclusion

We demonstrated that musical sophistication is associated with SPIN perception with a small effect size in a large population-based sample. Our results further showed that this relationship is fully mediated by AWM ability, and that the mediation effect remained significant in additional control analyses controlling for non-verbal intelligence, hearing thresholds, and age. In contrast, VWM did not explain this link. Non-verbal intelligence partially mediated the association and should therefore be considered a potential confound in future studies. Overall, our findings indicate that greater musical sophistication is associated with better SPIN perception in daily life, and that superior AWM ability appears to be key factor driving this advantage.

## References

Albert, D. (2006). Socioeconomic Status and Instrumental Music: What Does the Research Say about the Relationship and Its Implications? Update: Applications of Research in Music Education, 25, 39–45. 10.1177/87551233060250010105

Başkent, D., & Gaudrain, E. (2016). Musician advantage for speech-on-speech perception. J Acoust Soc Am, 139(3), El51-56. 10.1121/1.4942628

Bates, T. C., Lewis, G. J., & Weiss, A. (2013). Childhood socioeconomic status amplifies genetic effects on adult intelligence. Psychol Sci, 24(10), 2111–2116. 10.1177/0956797613488394

Benzaquén, E., Çolak, H., Guo, X., Lad, M., Rushton, S. P., & Griffiths, T. D. (2025). Auditory-cognitive contributions to speech-in-noise perception determined with structural equation modelling of a large sample. Sci Rep, 15(1), 34915. 10.1038/s41598-025-18800-6

Bidelman, G. M., & Yoo, J. (2020). Musicians Show Improved Speech Segregation in Competitive, Multi-Talker Cocktail Party Scenarios. Front Psychol, 11, 1927. 10.3389/fpsyg.2020.01927

Boebinger, D., Evans, S., Rosen, S., Lima, C. F., Manly, T., & Scott, S. K. (2015). Musicians and non-musicians are equally adept at perceiving masked speech. J Acoust Soc Am, 137(1), 378–387. 10.1121/1.4904537

Chierchia, G., Fuhrmann, D., Knoll, L. J., Pi-Sunyer, B. P., Sakhardande, A. L., & Blakemore, S. J. (2019). The matrix reasoning item bank (MaRs-IB): novel, open-access abstract reasoning items for adolescents and adults. R Soc Open Sci, 6(10), 190232. 10.1098/rsos.190232

Dryden, A., Allen, H. A., Henshaw, H., & Heinrich, A. (2017). The Association Between Cognitive Performance and Speech-in-Noise Perception for Adult Listeners: A Systematic Literature Review and Meta-Analysis. Trends in Hearing, 21, 2331216517744675. 10.1177/2331216517744675

Dubinsky, E., Wood, E. A., Nespoli, G., & Russo, F. A. (2019). Short-Term Choir Singing Supports Speech-in-Noise Perception and Neural Pitch Strength in Older Adults With Age-Related Hearing Loss. Front Neurosci, 13, 1153. 10.3389/fnins.2019.01153

Füllgrabe, C., & Rosen, S. (2016). On The (Un)importance of Working Memory in Speech-in-Noise Processing for Listeners with Normal Hearing Thresholds. Front Psychol, 7, 1268. 10.3389/fpsyg.2016.01268

Grassi, M., Talamini, F., Altoè, G., Brattico, E., Caclin, A., Carretti, B., Drai-Zerbib, V., Ferreri, L., Gambarota, F., Grahn, J., Guiotto Nai Fovino, L., Roccato, M., Rodriguez-Fornells, A., Swaminathan, S., Tillmann, B., Vuust, P., Wilbiks, J., Zentner, M., Aguilar, K.,…Zappa, A. (2025). Do Musicians Have Better Short-Term Memory Than Nonmusicians? A Multilab Study. Advances in Methods and Practices in Psychological Science, 8(4), 25152459251379432. 10.1177/25152459251379432

Guo, X., Benzaquén, E., Holmes, E., Choi, I., McMurray, B., Bamiou, D. E., Berger, J. I., & Griffiths, T. D. (2024). British version of the Iowa test of consonant perception. JASA Express Lett, 4(12). 10.1121/10.0034738

Hennessy, S., Mack, W. J., & Habibi, A. (2022). Speech-in-noise perception in musicians and non-musicians: A multi-level meta-analysis. Hear Res, 416, 108442. 10.1016/j.heares.2022.108442

Hester, R. L., Kinsella, G. J., & Ong, B. (2004). Effect of age on forward and backward span tasks. J Int Neuropsychol Soc, 10(4), 475–481. 10.1017/s1355617704104037

Holmes, E., & Griffiths, T. D. (2019). ’Normal’ hearing thresholds and fundamental auditory grouping processes predict difficulties with speech-in-noise perception. Sci Rep, 9(1), 16771. 10.1038/s41598-019-53353-5

Holster, J. D. (2023). The influence of socioeconomic status, parents, peers, psychological needs, and task values on middle school student motivation for school music ensemble participation. Psychology of Music, 51(2), 447–462. 10.1177/03057356221098095

Jahanshahi, M., Saleem, T., Ho, A. K., Fuller, R., & Dirnberger, G. (2008). A preliminary investigation of the running digit span as a test of working memory. Behav Neurol, 20(1-2), 17–25. 10.3233/ben-2008-0212

Jamey, K., Foster, N. E. V., Hyde, K. L., & Dalla Bella, S. (2024). Does music training improve inhibition control in children? A systematic review and meta-analysis. Cognition, 252, 105913. 10.1016/j.cognition.2024.105913

Kaplan, E. C., Wagner, A. E., Toffanin, P., & Başkent, D. (2021). Do Musicians and Non-musicians Differ in Speech-on-Speech Processing? Front Psychol, 12, 623787. 10.3389/fpsyg.2021.623787

Kim, S., Choi, I., Schwalje, A. T., Kim, K., & Lee, J. H. (2020). Auditory Working Memory Explains Variance in Speech Recognition in Older Listeners Under Adverse Listening Conditions. Clin Interv Aging, 15, 395–406. 10.2147/cia.S241976

Kraus, N., Strait, D. L., & Parbery-Clark, A. (2012). Cognitive factors shape brain networks for auditory skills: spotlight on auditory working memory. Ann N Y Acad Sci, 1252(1), 100–107. 10.1111/j.1749-6632.2012.06463.x

Lad, M., Billig, A. J., Kumar, S., & Griffiths, T. D. (2022). A specific relationship between musical sophistication and auditory working memory. Sci Rep, 12(1), 3517. 10.1038/s41598-022-07568-8

Lad, M., Holmes, E., Chu, A., & Griffiths, T. D. (2020). Speech-in-noise detection is related to auditory working memory precision for frequency. Sci Rep, 10(1), 13997. 10.1038/s41598-020-70952-9

Liu, M., Arseneau-Bruneau, I., Farrés Franch, M., Latorre, M. E., Samuels, J., Issa, E., Payumo, A., Rahman, N., Loureiro, N., Leung, T. C. M., Nave, K. M., von Handorf, K. M., Hoddinott, J. D., Coffey, E. B. J., Grahn, J., & Zatorre, R. J. (2025). Auditory working memory mechanisms mediating the relationship between musicianship and auditory stream segregation. Front Psychol, 16, 1538511. 10.3389/fpsyg.2025.1538511

Madsen, S. M. K., Marschall, M., Dau, T., & Oxenham, A. J. (2019). Speech perception is similar for musicians and non-musicians across a wide range of conditions. Scientific Reports, 9(1), 10404. 10.1038/s41598-019-46728-1

Madsen, S. M. K., Whiteford, K. L., & Oxenham, A. J. (2017). Musicians do not benefit from differences in fundamental frequency when listening to speech in competing speech backgrounds. Scientific Reports, 7(1), 12624. 10.1038/s41598-017-12937-9

Maillard, E., Joyal, M., Murray, M. M., & Tremblay, P. (2023). Are musical activities associated with enhanced speech perception in noise in adults? A systematic review and meta-analysis. Curr Res Neurobiol, 4, 100083. 10.1016/j.crneur.2023.100083

Maillard, E., Joyal, M., Murray, M. M., & Tremblay, P. (2023). Are musical activities associated with enhanced speech perception in noise in adults? A systematic review and meta-analysis. Current Research in Neurobiology, 4, 100083. 10.1016/j.crneur.2023.100083

Marsja, E., Holmer, E., Stenbäck, V., Micula, A., Tirado, C., Danielsson, H., & Rönnberg, J. (2025). Fluid Intelligence Partially Mediates the Effect of Working Memory on Speech Recognition in Noise. J Speech Lang Hear Res, 68(1), 399–410. 10.1044/2024_jslhr-24-00465

Merten, N., Boenniger, M. M., Herholz, S. C., & Breteler, M. M. B. (2022). The Associations of Hearing Sensitivity and Different Cognitive Functions with Perception of Speech-in-Noise. Ear Hear, 43(3), 984–992. 10.1097/aud.0000000000001154

Moberly, A. C., Vasil, K. J., Wucinich, T. L., Safdar, N., Boyce, L., Roup, C., Holt, R. F., Adunka, O. F., Castellanos, I., Shafiro, V., Houston, D. M., & Pisoni, D. B. (2018). How does aging affect recognition of spectrally degraded speech? Laryngoscope, 128 Suppl 5(Suppl 5). 10.1002/lary.27457

Moore, D. R., Edmondson-Jones, M., Dawes, P., Fortnum, H., McCormack, A., Pierzycki, R. H., & Munro, K. J. (2014). Relation between speech-in-noise threshold, hearing loss and cognition from 40-69 years of age. PLoS One, 9(9), e107720. 10.1371/journal.pone.0107720

Morse-Fortier, C., Parrish, M. M., Baran, J. A., & Freyman, R. L. (2017). The Effects of Musical Training on Speech Detection in the Presence of Informational and Energetic Masking. Trends in Hearing, 21, 2331216517739427. 10.1177/2331216517739427

Müllensiefen, D., Gingras, B., Musil, J., & Stewart, L. (2014). The musicality of non-musicians: an index for assessing musical sophistication in the general population. PLoS One, 9(2), e89642. 10.1371/journal.pone.0089642

Parbery-Clark, A., Skoe, E., Lam, C., & Kraus, N. (2009). Musician enhancement for speech-in-noise. Ear Hear, 30(6), 653–661. 10.1097/AUD.0b013e3181b412e9

Parbery-Clark, A., Strait, D. L., Anderson, S., Hittner, E., & Kraus, N. (2011). Musical experience and the aging auditory system: implications for cognitive abilities and hearing speech in noise. PLoS One, 6(5), e18082. 10.1371/journal.pone.0018082

Patel, A. D. (2011). Why would Musical Training Benefit the Neural Encoding of Speech? The OPERA Hypothesis. Front Psychol, 2, 142. 10.3389/fpsyg.2011.00142

Patel, A. D. (2014). Can nonlinguistic musical training change the way the brain processes speech? The expanded OPERA hypothesis. Hear Res, 308, 98–108. 10.1016/j.heares.2013.08.011

Román-Caballero, R., Vadillo, M. A., Trainor, L. J., & Lupiáñez, J. (2022). Please don’t stop the music: A meta-analysis of the cognitive and academic benefits of instrumental musical training in childhood and adolescence. Educational Research Review, 35, 100436. 10.1016/j.edurev.2022.100436

Rönnberg, J., Lunner, T., Zekveld, A., Sörqvist, P., Danielsson, H., Lyxell, B., Dahlström, O., Signoret, C., Stenfelt, S., Pichora-Fuller, M. K., & Rudner, M. (2013). The Ease of Language Understanding (ELU) model: theoretical, empirical, and clinical advances. Front Syst Neurosci, 7, 31. 10.3389/fnsys.2013.00031

Rosenthal, E. N., Riccio, C. A., Gsanger, K. M., & Jarratt, K. P. (2006). Digit Span components as predictors of attention problems and executive functioning in children. Archives of Clinical Neuropsychology, 21(2), 131–139. 10.1016/j.acn.2005.08.004

Ruggles, D. R., Freyman, R. L., & Oxenham, A. J. (2014). Influence of musical training on understanding voiced and whispered speech in noise. PLoS One, 9(1), e86980. 10.1371/journal.pone.0086980

Schellenberg, E. (2011). Music Lessons, Emotional Intelligence, and IQ. Music Perception, 29, 185–194. 10.1525/mp.2011.29.2.185

Schellenberg, E. G., & Lima, C. F. (2024). Music Training and Nonmusical Abilities. Annu Rev Psychol, 75, 87–128. 10.1146/annurev-psych-032323-051354

Schulze, K., & Koelsch, S. (2012). Working memory for speech and music. Ann N Y Acad Sci, 1252, 229–236. 10.1111/j.1749-6632.2012.06447.x

Slater, J., Skoe, E., Strait, D. L., O’Connell, S., Thompson, E., & Kraus, N. (2015). Music training improves speech-in-noise perception: Longitudinal evidence from a community-based music program. Behav Brain Res, 291, 244–252. 10.1016/j.bbr.2015.05.026

Smith, S. B., Krizman, J., Liu, C., White-Schwoch, T., Nicol, T., & Kraus, N. (2019). Investigating peripheral sources of speech-in-noise variability in listeners with normal audiograms. Hear Res, 371, 66–74. 10.1016/j.heares.2018.11.008

Stam, M., Smit, J. H., Twisk, J. W., Lemke, U., Smits, C., Festen, J. M., & Kramer, S. E. (2016). Change in Psychosocial Health Status Over 5 Years in Relation to Adults’ Hearing Ability in Noise. Ear Hear, 37(6), 680–689. 10.1097/aud.0000000000000332

Stevenson, J. S., Clifton, L., Kuźma, E., & Littlejohns, T. J. (2022). Speech-in-noise hearing impairment is associated with an increased risk of incident dementia in 82,039 UK Biobank participants. Alzheimers Dement, 18(3), 445–456. 10.1002/alz.12416

Talamini, F., Altoè, G., Carretti, B., & Grassi, M. (2017). Musicians have better memory than nonmusicians: A meta-analysis. PLoS One, 12(10), e0186773. 10.1371/journal.pone.0186773

von Stumm, S., & Plomin, R. (2015). Socioeconomic status and the growth of intelligence from infancy through adolescence. Intelligence, 48, 30–36. 10.1016/j.intell.2014.10.002

Yeend, I., Beach, E. F., & Sharma, M. (2019). Working Memory and Extended High-Frequency Hearing in Adults: Diagnostic Predictors of Speech-in-Noise Perception. Ear Hear, 40(3), 458–467. 10.1097/aud.0000000000000640

Yoo, J., & Bidelman, G. M. (2019). Linguistic, perceptual, and cognitive factors underlying musicians’ benefits in noise-degraded speech perception. Hear Res, 377, 189–195. 10.1016/j.heares.2019.03.021

Yurgil, K. A., Velasquez, M. A., Winston, J. L., Reichman, N. B., & Colombo, P. J. (2020). Music Training, Working Memory, and Neural Oscillations: A Review [Review]. Frontiers in Psychology, Volume 11 - 2020. 10.3389/fpsyg.2020.00266

Zhang, L., Fu, X., Luo, D., Xing, L., & Du, Y. (2021). Musical Experience Offsets Age-Related Decline in Understanding Speech-in-Noise: Type of Training Does Not Matter, Working Memory Is the Key. Ear Hear, 42(2), 258–270. 10.1097/aud.0000000000000921

